# CMPyMOL: An Interactive PyMOL extension for Protein Contact-Map Analysis

**DOI:** 10.1101/084269

**Authors:** Venkatramanan Krishnamani

**Affiliations:** Department of Molecular Physiology and Biophysics, University of Iowa, Iowa 52242, US

**Keywords:** contact maps, pymol, biochemical, structure

## Abstract

Contact–maps are reduced 2D representation of the 3D spatial configuration of a protein. Many valuable structural features like secondary structures, inter- and intra-protein interactions,interacting domains, etc., can be readily identified from these maps. However, it is not straightforward and intuitive to reckon the spatial organization of the contact regions from reduced representation. The CMPyMOL extention for molecular visualization software PyMOL attempts to bridge this gap as an interactive graphical tool for protein contact-maps that interfaces with PyMOL for 3D visualization. Specifically, CMPyMOL helps understand the functional importance of contacts by providing visual overlays of various structural and biochemical properties of a protein on top of its contact-map.

## 1. Introduction

A contact-map of a protein is a 2D matrix of pairwise inter-residue distances, typically calculated over distances between C*α* atoms subject to an arbitrary maximum threshold. By construction this distance matrix is square and symmetrical. Contact-maps have been traditionally used to compare two protein structures/conformations [1], protein-protein-interactions [2, 3], protein folding [4, 5], structure prediction [6] and even reconstruction of the protein’s 3D structure [7].

## 2. Problems and Background

Contact-maps capture high resolution, residue-level information regarding local conformations such as *α*-helices and *β*-sheets, and non-local interactions like inter-domain interactions. In this respect, contact-maps are a loss-less representation of structural information (except for chirality). However,essential biochemical information such as the residue type and the properties associated with it are lost during a contact-maps’ construction. The interac-tion type of a particular contact point, such as hydrophobic interactions, salt bridges and hydrogen bonds, etc., can be crucial in understanding protein structure and function. A manual assignment to keep track of the residue number, the residue-type from the protein sequence and its spatial location can quickly become cumbersome and unmanageable.

## 3. Software Architecture

CMPyMOL supplements a contact-map analysis by interfacing with the powerful 3D visualization capabilities of PyMOL [8]. Launching CMPyMOL automatically invokes the PyMOL executable and generates a contact-map (for a specified cut-off distance) for a given PDB file. Visualizing multi-frame PDB trajectories are also supported. The user is provided with an option to calculate the distance map between C*α* or C*β* atoms of residue pairs and the desired cutoff distance. The main program window (**Figure 1**) displays the contact-map in gray scale, shaded according to the distance between the pair of C*α* atoms.

**Figure 1:**
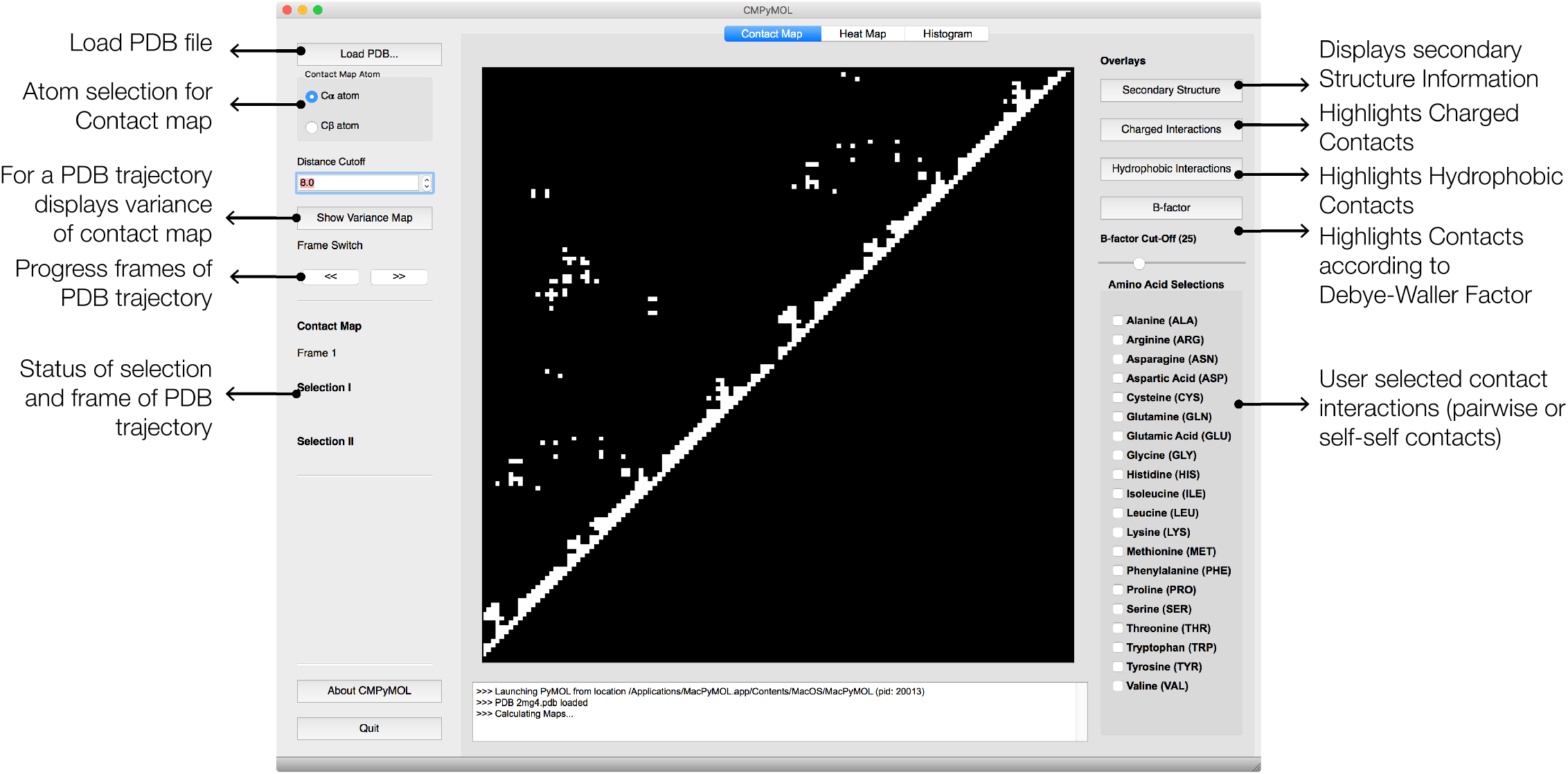
The main window of CMPyMOL provides controls for all the selection,overlay and plots to analyze contact maps. The overlays (the toggle buttons onthe right of the contact map) superpose chemical and structural information on top of the contact map when activated. The plots (buttons on the left side of the contact map) pops open a new window that provides an overview of the nature of contacts.

CMPyMOL allows for manual selection of interacting residues on the 2D contact map while the program highlights the corresponding residues in the PyMOL 3D visualization. This provides a intuitive bridge between the 2D and 3D representations of the protein. It also provides visual over-lays of structural and biochemical properties of the amino acid residues.Secondary structure (using STRIDE [9, 10] embedded within CMPyMOL),charge-charge interactions, hydrophobic interactions, B-factor and custom selected residue pair interaction sites (see **Figure 1**) are the currently available overlays in CMPyMOL. The program also calculates pairwise residue interaction heat-map and residue-wise contact density plots as an alternate representations of the contact-map data.

## 4. Highlighted Software Functionalities

### 4.1. Substructure Selection

Contact points can be selected individually (left-click) or as contact regions (rectangular selection by left-click and drag) interactively over the contact-map. When a substructure of interest is selected, the corresponding residues are synchronously highlighted in the PyMOL window (**Figure 2A and 2B**). The secondary structure overlay shows that our selection is a contact formed between two helical segments (*red* rectangles in **Figure 2C**). Since the example protein is a homodimeric protein (PDBID: 2MG4) [11], the gray square represents the inter–protein interaction region and the selection corresponds to the monomer–monomer contact.

It should be noted that CMPyMOL only highlights the residues that are in contact and within the current selection. By construction, the lower and upper half of the contact-map separated by the main diagonal are symmetric, hence the lower half of the contact map is not shown by default.

**Figure 2:**
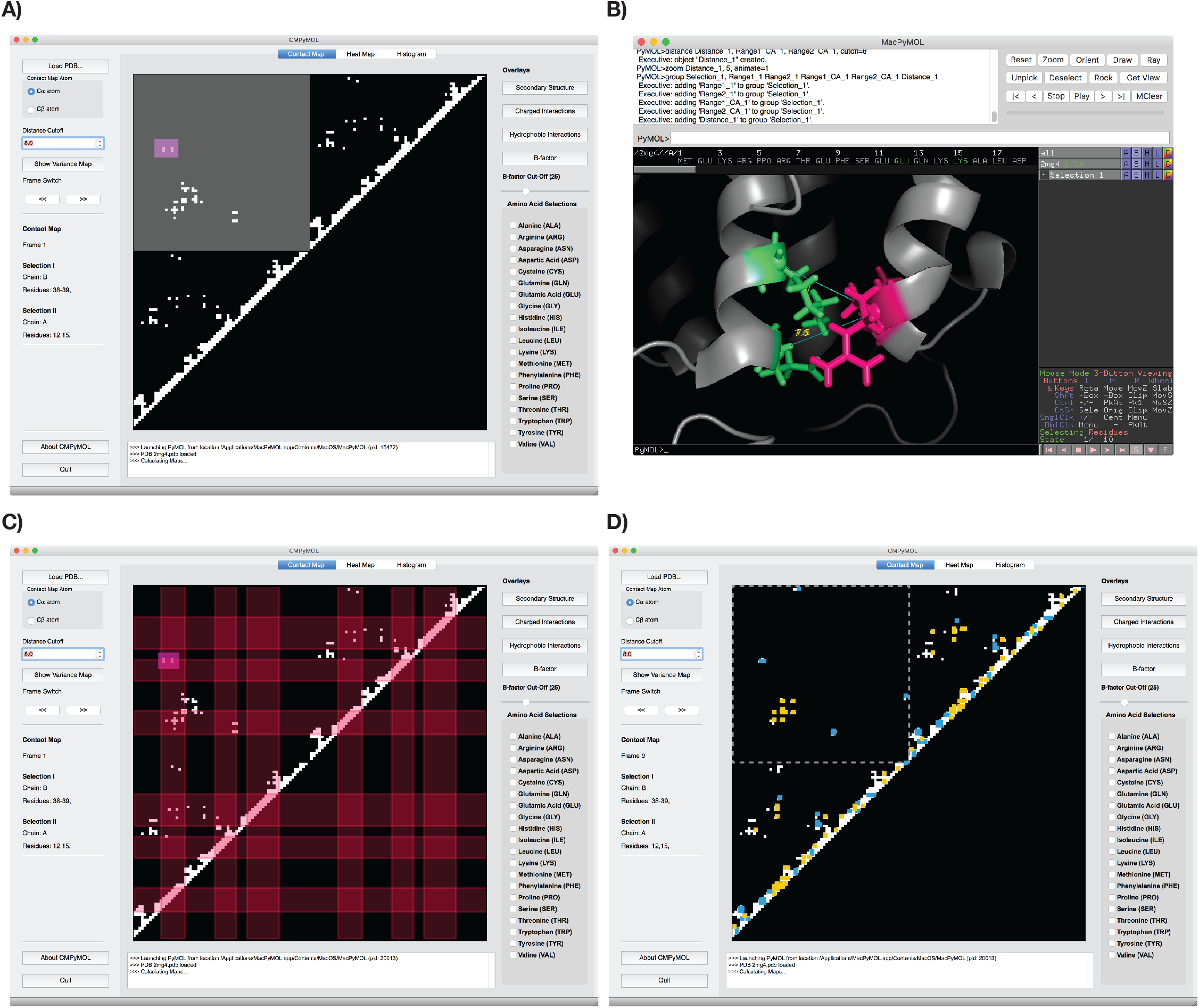
**A)** The purple rectangular selection highlights a contact in the inter–protein interaction region (gray square) of the contact map. The contact map was generated with a cutoff of 8Å between C*α* atoms of PDB:2MG4. **B)** This selection also brings to focus the selected region in the PyMOL window. The interacting residues within the selection are highlighted in red and blue with the sidechains represented as sticks. **C)** Activating the secondary structure overlay identifies the previous selection is a contact between two *α*-helices. **D)** To overlay the chemical nature, namely, charged (blue) and hydrophobic (yellow), highlights the corresponding contacts.

### 4.2. Overlays and Plots

A novel feature in CMPyMOL are the overlays of structural and biochemical properties that highlights the nature of interaction of each contact point. This allows clear distinction, discovery and classification of contacts of a specific biochemical/structural type. For example, turning on “Charged Interactions” displays a *blue* overlay on all the contacts points where two charged residues are in proximity, i.e. in contact (while “Hydrophobic Interactions” as displayed as an *yellow* overlay) (**Figure 2D**). “B-factor” draws a *red* highlight of residues that have a Debye-Waller factor [12, 13] above a certain cutoff (specified by the slider, **Figure 1**) and that are in contact (*not shown*). Additionally, users can select any pair of amino-acids, listed on the right, to highlight those residues selections in PyMOL window that are within the defined cutoff (*see manual*). This is a powerful tool for quickly and efficiently identifying specific interactions types and simultaneously visualizing their spatial orientation.

The “Pairwise Heat-map” and “Contacts Histogram” (**Figure 3**), calculates and plots the number of pairwise residue contacts and contact density of each residue, respectively.Of these maps, the contact density map is interactive, mouse selections of a particular contact density highlights the location in the PyMOL window. When a multiframe PDB trajectory is loaded into the software, the user can choose to view either the contact-map or the cumulative variance contact-map calculated for all the frames starting from frame 1 to the current frame selection.

**Figure 3:**
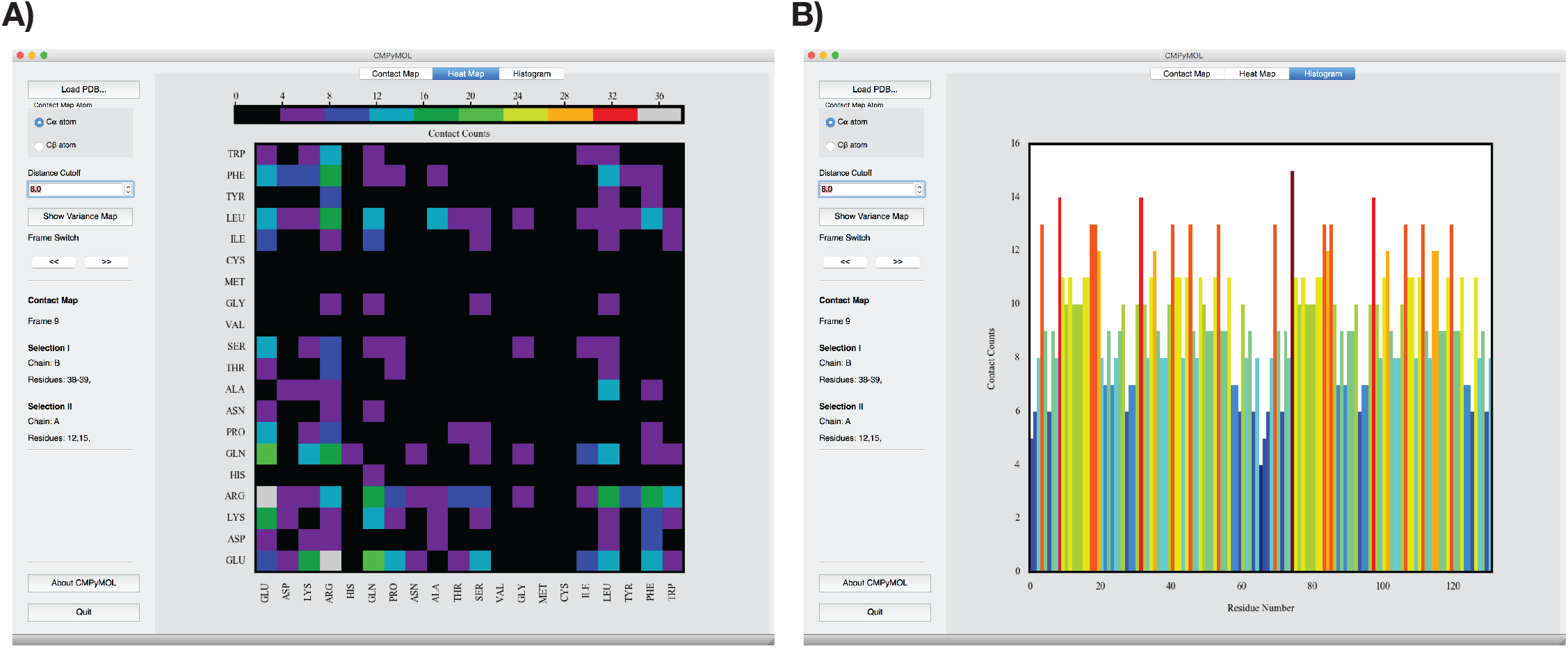
**A)** Heatmap of pairwise residue-residue interaction map. This map counts the number of pairwise contacts of a given aminoacid to the rest of the other aminoacids. The order of aminoacids in this plot are arranged according to their hydrophobicity. The color scale shows the number of each pairwise contacts in the protein. **B)** The residue-wise density of contacts in the protein (PDBID:2MG4). The contact histogram plot, graphs the density of contacts with respect to residue position. Both plots are interactive and clicking on a particular residue or residue-pair highlights the corresponding selection in the PyMOL window.

## 5. Implementation

CMPyMOL is developed using the Python programming language and open source libraries–*PyQT4*, Numeric Python (*numpy*) and *matplotlib*. It is provided under an open source license (The MIT License) as source code and as pre-compiled binaries for Windows and Mac OS X operating systems (Linux users will be able to run CMPyMOL directly from source). The detailed installation instructions are listed in the user guide (**Table 1**).

**Table 1:** Software metadata.

|  |  |  |
| --- | --- | --- |
| S1 | Current software version | 2.0 |
| S2 | Permanent link to executables of this version | https://github.com/emptyewer/CMPyMOL/releases |
| S3 | Legal Software License | MIT License |
| S4 | Computing platform/Operating System | Windows and Mac OS X |
| S5 | Installation requirements & dependencies | *pymol* |
| S6 | User manual | https://github.com/emptyewer/CMPyMOL/releases |
| S7 | Support email for questions |  |

CMPyMOL provides a much needed add-on to the PyMOL software package, a tool which is typically a built-in part of other molecular visualization programs, such as VMD [14]. There exists at least one other interactive tool for visualizing contact-maps, but it is limited in terms of displaying biochemical and structural information on the contact-maps and does not support multi-frame PDB trajectories [15]. CMPyMOL is intended to replace an existing PyMOL plugin, Contact Map Visualizer (available from *http://www.pymolwiki.org*), that was co-developed by the author (VK) in collaboration with Thomas Holder (Schro¨dinger, LLC).

## 6. Illustrative Example

Using an NMR structure (10 models) of a designed homodimeric protein (PDBID: 2MG4), one workflow highlighting three core functionalities of CMPyMOL will be demonstrated in this section [11]. CMPyMOL user guide is downloadable from the software’s github page.It provides detailed descriptions of other functionality of CMPyMOL. https://github.com/emptyewer/CMPyMOL. The PDB file (PDBID: 2MG4) used in this example is distributed along with the source code.

### 6.1. Identifying Chemical Nature of Secondary Structure Contacts

After the CMPyMOL window is initialized and the PDB file is loaded, selecting the toggle button (named Secondary Structure) on the right–hand side of the contact map main–display overlays the secondary structure information. The secondary structure overlay superposes *α*-helical and *β*-sheet as red and green translucent rectangles, respectively (**Figure 2C**). The loop regions are uncolored. Selecting the contact points within the *α*-helix–*α*-helix interaction region brings to focus in the corresponding 3D structure in the Py-MOL window (**Figure 2B**). Further examining the selection in the PyMOL window reveals the biochemistry of the interaction is electorstatic/charged in nature (see *blue* squares in **Figure 2D** corresponding to the selection where GLU, LYS and ARG within 8Å).

### 6.2. Identifying Contacts that Stabilizes Protein Dimer

In **Figure 2A** the gray box on the top left represents the contacts from inter–protein monomer–monomer interaction. The secondary structure overlay readily identifies most of the contact between the protein partners are *α*-helix–*α*-helix interaction (contact points within the region where two red rectangles intersect) with some interactions between two loops (contacts within uncolored region) and a few interactions between *α*-helix and loop **Figure 2C**. Further the interactions stabilizing the dimer are hydrophobic (*yellow* squares within *dotted-gray* box **Figure 2D**) and *two* charged interactions (*blue* squares **Figure 2D**). Note that even though there are four *blue* spots within the *gray* box, due to diagonal symmetry of the contact map the actual charged interactions in the protein is two.

### 6.3. Identifying Regions of Maximum Flexibility

Since the PDB in this example is an NMR structure, CMPyMOL can calculate the variance of the distances between pairwise contacts along the trajectory. With such a representation, selecting a region displaying highest variance in intra–protein interaction region reveals that the contacts belong to a particular loop on each monomer (*not shown*). This is described in more detail in the CMPyMOL user manual that can be downloaded from the offical site. https://github.com/emptyewer/CMPyMOL/releases

## 7. Conclusions and Limitations

CMPyMOL integrates 2D contact-maps augmented with biochemical information and powerful 3D Visualization of PyMOL. This provides an intuitive platform for simultaneously exploring protein interaction sites and its 3D structure.

Currently, CMPyMOL only supports importing locally available PDB file-formatted files. Since each pixel of the contact-map image represents an interacting residue pair, the number of residues of a protein that can be comfortably displayed on a computer screen is limited to approximately 1000px by 1000px or 1500px by 1500px (varies by screen resolution). This software is under active development, so users can request new features and report bugs on the CMPyMOL github repository. The next release (2.1) will support further data formats.

## Acknowledgements

I would like to thank my former postdoctoral advisor Dr. Markus Deserno at Carnegie Mellon University for his support and insightful suggestions.*Funding*: Volkwagen Foundation for supporting this project within the framework of their program “New Conceptual Approaches to Modeling and Simulation of Complex Systems”

